# Ancient MAPK ERK7 is regulated by an unusual inhibitory scaffold required for Toxoplasma apical complex biogenesis

**DOI:** 10.1101/2020.02.02.931089

**Authors:** Peter S. Back, William J. O’Shaughnessy, Andy S. Moon, Pravin S. Dewangan, Xiaoyu Hu, Jihui Sha, James A. Wohlschlegel, Peter J. Bradley, Michael L. Reese

**Affiliations:** Molecular Biology Institute, University of California, Los Angeles, Los Angeles, California; Department of Pharmacology, University of Texas, Southwestern Medical Center, Dallas, TX USA; Department of Molecular Microbiology and Immunology, University of California, Los Angeles, CA USA; Department of Biological Chemistry, David Geffen School of Medicine, University of California, Los Angeles, Los Angeles, California, USA; Department of Biochemistry, University of Texas, Southwestern Medical Center, Dallas, TX USA

**Keywords:** Kinase, scaffold, intrinsically-disordered protein, cilium

## Abstract

Apicomplexan parasites use a specialized cilium structure called the apical complex to organize their secretory organelles and invasion machinery. The apical complex is integrally associated with both the parasite plasma membrane and an intermediate filament cytoskeleton called the inner membrane complex (IMC). While the apical complex is essential to the parasitic lifestyle, little is known about the regulation of apical complex biogenesis. Here, we identify AC9 (apical cap protein 9), a largely intrinsically disordered component of the *Toxoplasma gondii* IMC, as essential for apical complex development, and therefore for host cell invasion and egress. Parasites lacking AC9 fail to successfully assemble the tubulin-rich core of their apical complex, called the conoid. We use proximity biotinylation to identify the AC9 interaction network, which includes the kinase ERK7. Like AC9, ERK7 is required for apical complex biogenesis. We demonstrate that AC9 directly binds ERK7 through a conserved C-terminal motif and that this interaction is essential for ERK7 localization and function at the apical cap. The crystal structure of the ERK7:AC9 complex reveals that AC9 is not only a scaffold, but also inhibits ERK7 through an unusual set of contacts that displaces nucleotide from the kinase active site. ERK7 is an ancient and auto-activating member of the mitogen-activated kinase family and we have identified its first regulator in any organism. We propose that AC9 dually regulates ERK7 by scaffolding and concentrating it at its site of action while maintaining it in an “off” state until the specific binding of a true substrate.

**Significance Statement:** Apicomplexan parasites include the organisms that cause widespread and devastating human diseases such as malaria, cryptosporidiosis, and toxoplasmosis. These parasites are named for a structure, called the “apical complex,” that organizes their invasion and secretory machinery. We found that two proteins, apical cap protein 9 (AC9) and an enzyme called ERK7 work together to facilitate apical complex assembly. Intriguingly, ERK7 is an ancient molecule that is found throughout Eukaryota, though its regulation and function are poorly understood. AC9 is a scaffold that concentrates ERK7 at the base of the developing apical complex. In addition, AC9 binding likely confers substrate selectivity upon ERK7. This simple competitive regulatory model may be a powerful but largely overlooked mechanism throughout biology.

## Introduction

Cilia are ancient eukaryotic organelles that organize signal transduction cascades and mediate cell motility. These functions are driven by the cooperation of cytoskeleton and membrane structures and require specialized signaling and trafficking machinery for their biogenesis and maintenance (1–3). In apicomplexan parasites, the cilium is thought to have evolved to form the “apical complex,” (4–7) which organizes the parasites’ invasion machinery and for which the phylum is named. Apicomplexa include the causative agents of malaria, toxoplasmosis, and cryptosporidiosis, which all invade mammalian cells to cause disease. Like more typical eukaryotic cilia, the apical complex is composed of specialized microtubule structures inserted into the plasma membrane (8). In addition, the apical complex is the site of secretion of specialized organelles called micronemes and rhoptries that mediate the parasites’ attachment to and invasion of host cells. In the asexual stage of most apicomplexans, secretion is thought to occur through a tubulin-rich structure in the apical complex called the conoid (8, 9). The apical complex is also intimately associated with an intermediate filament cytoskeleton called the inner membrane complex (IMC) that scaffolds the apicomplexan cell, ensuring its correct morphology. The IMC anchors the parasite actin-based motility machinery (10), powering the parasite’s motility as it glides across and invades host cells. While the IMC extends the length of the parasite, it has clearly segregated apical, medial, and basal sub-domains that are defined by specific protein localization (11, 12). In *Toxoplasma gondii*, the IMC “apical cap” comprises the anterior-most ∼1 μm of the IMC, just basal to the apical complex. The apical cap appears to be a site at which actin regulators (13) and subcomponents of the parasite invasion machinery concentrate (14). While a number of IMC proteins have been genetically manipulated and evaluated phenotypically, their biochemical functions have been understudied.

In the present work, we identify one component of the *Toxoplasma gondii* apical IMC, apical cap protein 9 (AC9), as essential to the parasite lytic cycle. We found that loss of AC9 results in parasites that are entirely unable to egress from their host cells or invade new cells. These deficiencies are attributable to the failure of the parasites to form a functional apical complex, as the conoids are entirely missing in mature parasites and regulated secretion is disrupted. These data provide new insight into the functions of the IMC apical cap in regulating parasite development. Using proximity biotinylation, we defined the AC9 interaction network, which includes ERK7, a conserved MAPK that regulates ciliogenesis in metazoa (15, 16), and which we have recently shown is required for conoid formation (17). We demonstrate that AC9 is required for the correct localization of ERK7 at the apical cap, and that this scaffolding interaction is essential for apical complex maturation. Finally, we solved the crystal structure of the ERK7:AC9 complex, which revealed that the AC9 C-terminus wraps around the kinase and inserts into the active site, inhibiting it. ERK7 orthologs are found in all eukaryotes with ciliated cells, though the pathways it regulates are largely unknown. Furthermore, ERK7 is autoactivating (18), and we have identified the first regulatory interaction for this ancient kinase in any organism. Moreover, our data highlight a simple competitive mechanism by which protein-protein interactions can ensure the fidelity and specificity of a signaling network.

## Results

### Loss of AC9 blocks host cell invasion and egress and parasite conoid assembly

AC9 (TGGT1_246950) was initially identified by proximity biotinylation as an apically localized interactor of the IMC suture component ISC4 (19), though AC9 function was not further investigated in the previous study. We were unable to obtain an AC9 knockout parasite strain using CRISPR/Cas9, thus we chose to assess its function using the auxin-inducible degron (AID) system (20, 21). We endogenously tagged the C-terminus of AC9 with an AID-3xHA tag (AC9^AID^). AC9^AID^ faithfully localized to the apical cap (Figure 1A) and addition of IAA (AC9^AID/IAA^) resulted in efficient degradation of the protein (Figure 1A, S1A). Loss of AC9 completely blocked the parasites’ ability to form plaques, which was rescued by the expression of a non-degradable copy of AC9 (Figure 1B). We found that AC9^AID/IAA^ parasites replicated normally but failed to egress from their host cells. Instead, AC9^AID/IAA^ parasites appeared to replicate until their vacuoles separated from the monolayer, which we found floating in the medium (Figure 1C). This phenotype is similar to the knockout of parasite perforin-like protein 1 (PLP1) (22), and suggested a block in egress. Egress can be induced by treatment with the calcium ionophore A23187 (23). While AC9^AID^ parasites efficiently egressed from host cells after ionophore treatment, AC9^AID/IAA^ were completely unresponsive (Figure 1D). Loss of AC9 also blocked parasite invasion of host cells (Figure 1E). Invasion and egress require secretion from specialized organelles called micronemes (24). The microneme protein MIC2 is shed from the parasite plasma membrane after secretion, and the levels of this protein in media can be used as a surrogate for secretion (23). Loss of AC9 blocked both basal and ionophore-induced release of MIC2, though levels of GRA39, which is constitutively secreted through a different route, are unaffected (Figure 1F). Taken together, our data show that AC9 is required for efficient microneme secretion, and its loss blocks the parasite lytic cycle.

**Figure 1:**
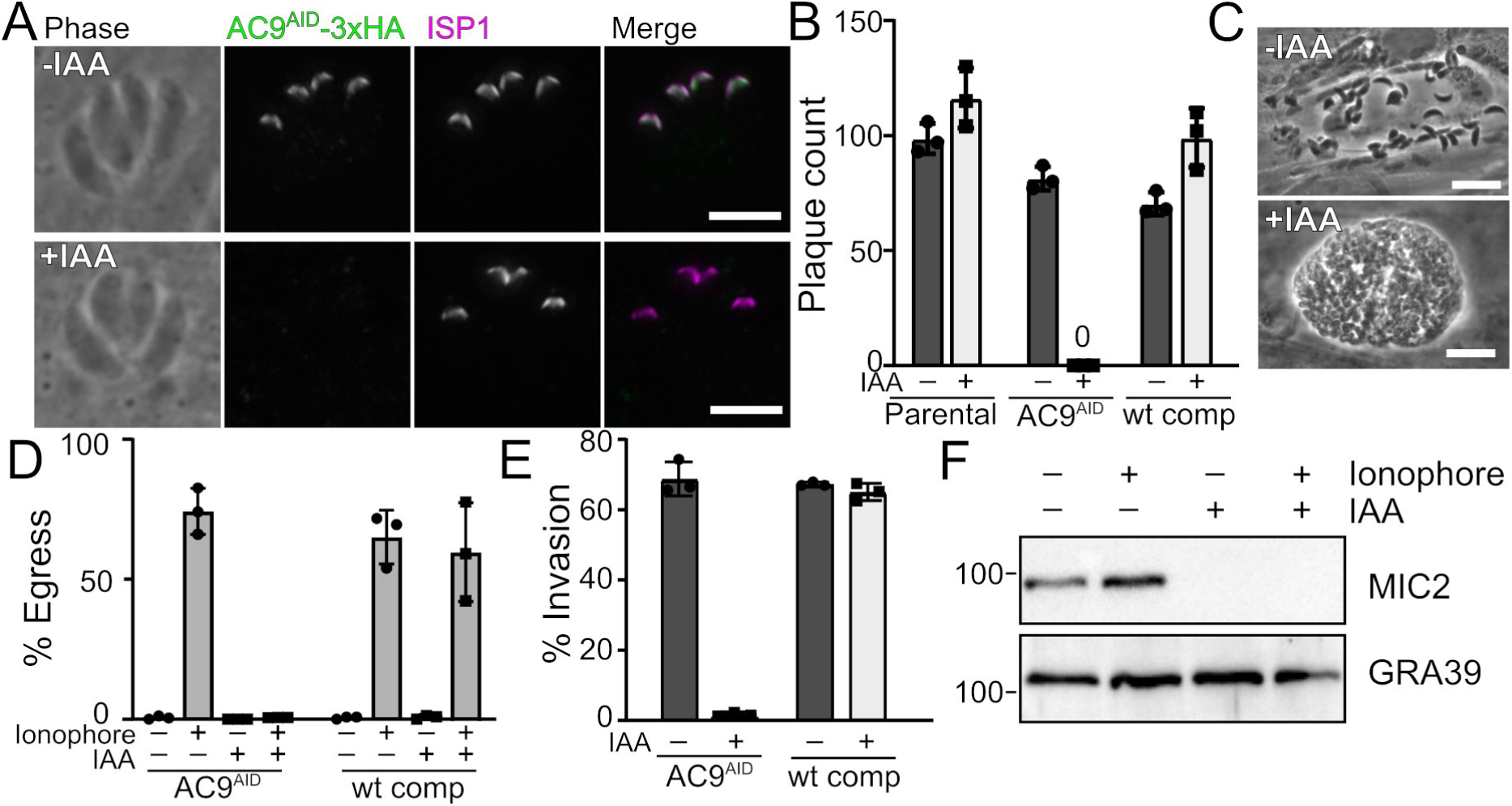
AC9 is required to complete the parasite lytic cycle. (A) AC9^AID^-3xHA (green) localizes to the apical cap (magenta) and is lost when parasites are treated with IAA. Scale bars: 5 μm. (B) Quantification of plaque number comparing growth of parental, AC9^AID^, and wt complemented AC9^AID^ parasites grown with and without IAA. (C) AC9^AID^ parasites naturally egress from host cells in -IAA but are found as floating vacuoles when grown in IAA. Scale bars: 20 μm. (D) Quantification of egress of the indicated strains induced by calcium ionophore and grown (-/+) IAA. (E) Quantification of invasion of the indicated strains grown (-/+) IAA. (F) Western blot of soluble secreted proteins from AC9^AID^ and AC9^AID/IAA^. Microneme secretion was tracked with anti-MIC2 and the constitutively secreted dense granule protein GRA39 was used as a control.

As AC9 impacts invasion and microneme secretion, we reasoned the observed phenotypes may be due to changes in parasite ultrastructure upon AC9 degradation. To test this hypothesis, we used immunofluorescence to compare available apical markers in AC9^AID^ and AC9^AID/IAA^ parasites. Strikingly, loss of AC9 resulted in a disorganization of the rhoptry secretory organelles (Figure 2A). These organelles are usually bundled and polarized with their necks pointing apically (marked by RON11). In AC9^AID/IAA^ parasites, however, the rhoptries are detached from the parasite’s apex and the necks are no longer consistently apically oriented (Figure 2A). Thus, both the micronemes and rhoptries are impacted in the absence of AC9. In *Toxoplasma*, a tubulin-rich structure called the conoid forms the core of the apical complex (8) and is the site at which microneme and rhoptry secretion is thought to occur (9, 25). Partial disruptions in the conoid structure have recently been associated with loss of parasite motility, invasion, and egress (26–28). Thus, we tested whether the conoid marker SAS6L (4) was altered upon AC9 degradation. In normally developed AC9^AID^ parasites, we observed distinct apical SAS6L foci in both mother and developing daughter parasites (Figure 2B). Strikingly, in AC9^AID/IAA^ parasites, the mother SAS6L signal was missing, while the daughter cells’ was maintained (Figure 2B). These data strongly suggest that AC9 is required for maturation of the parasite conoid, and that the invasion and egress phenotypes we observed upon AC9 degradation (Figure 1) are driven by conoid loss.

**Figure 2:**
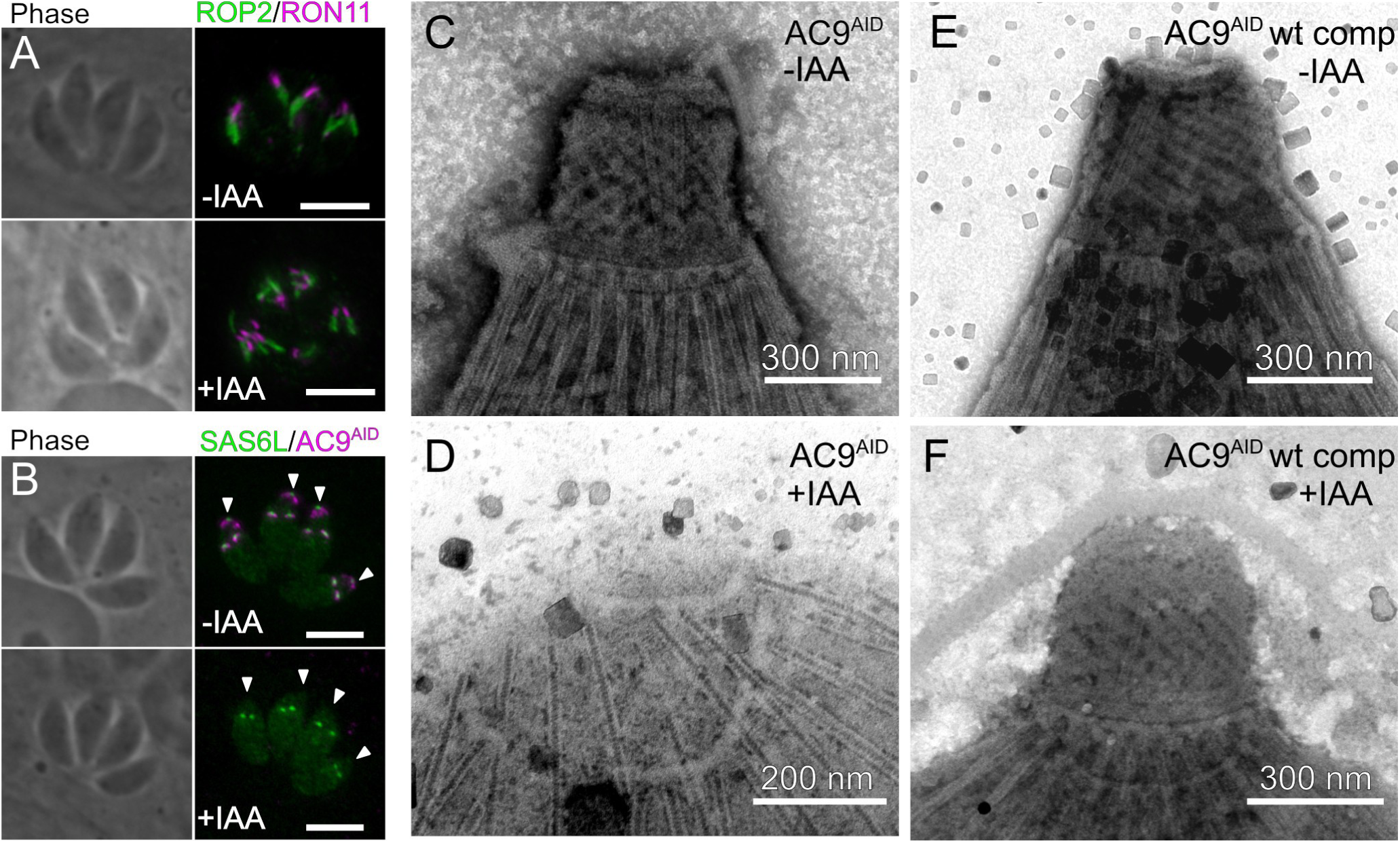
Loss of AC9 disrupts the parasite apical complex. (A) AC9^AID^ and AC9^AID/IAA^ parasites were stained with ROP2 (green) and RON11 (magenta). (B) AC9^AID^ and AC9^AID/IAA^ parasites were stained with antibodies to the HA tag (magenta) and the conoid marker SAS6L (green). Arrowheads indicate the position of the maternal apical complex. Scale bars: 5 μm. TEMs of the apical complex from negatively stained detergent-extracted (C) AC9^AID^ parasites, (D) AC9^AID/IAA^ parasites, and AC9^AID^ wt complement parasites grown in (E) -IAA and (F) +IAA.

We next sought to directly examine the effect of AC9 loss on the ultrastructure of the mature parasite conoid. Many parasite cytoskeletal structures, including the apical complex, are preserved after detergent extraction (29). We detergent extracted AC9^AID^ and AC9^AID/IAA^ parasites to create cytoskeleton “ghosts” and imaged them using negative stained transmission electron microscopy (TEM). While we observed an extended conoid in AC9^AID^ parasites (Figure 2C), we found that the conoid was completely absent in AC9^AID/IAA^ parasites (Figure 2D). As expected, expression of non-degradable AC9 rescued this phenotype (Figure 2E,F). Importantly, the ultrastructural changes appear to be specific to the loss of the conoid in AC9^AID/IAA^ parasites, as our TEMs show that the parasites have maintained their 22 cortical microtubules (Figure 2D). Intriguingly, we noted that AC9^AID^-tagging appears to have exacerbated an artifact of EM sample preparation of the parasite cytoskeleton in which the base of the conoid partially detaches from the apical polar ring (Figure S1). We assessed the basal (-IAA) level of AC9^AID^ protein, and found that AC9^AID^ levels are ∼40% of the levels of AC9 tagged with only a 3xHA in the same OsTIR1-expressing parental line (Figure S1). Such basal degradation has been uncovered as a common artifact of the AID system (30, 31). Notably, AC9^AID^(wt-comp) conoids were indistinguishable from the parental strain, suggesting that reduced AC9 protein levels may be driving the apparent defect in conoid ultrastructure (Figures 1, S1). Nevertheless, in spite of any potential structural differences, the AC9^AID^ parasites showed no phenotype in secretion, invasion, or egress when untreated with IAA, in stark contrast to the effect of complete AC9 degradation (Figure 1). Taken together, our data indicate AC9 has a critical role in maintaining apical complex ultrastructure.

### Proximity biotinylation reveals AC9 interacts with the MAP kinase ERK7

We next sought to identify protein partners that collaborate with AC9 to facilitate parasite conoid assembly and/or maintenance. To this end, we endogenously tagged the protein with BirA* (AC9^BioID^) (12, 32). After 24 h growth in 150 μM biotin, we could detect biotin labeling in the apical cap with fluorescent streptavidin, demonstrating active BirA* at this location (Figure S2A). While AC9 is predicted to be ∼70% intrinsically disordered, it has a ∼150 residue N-terminal predicted coiled-coil domain, which we reasoned may indicate it is a component of the IMC cytoskeleton. The IMC cytoskeleton is stable after detergent extraction (33), and we found that AC9 co-purified with cytoskeletal, rather than membranous, components of the IMC (Figure S2B). This demonstrates AC9 is associated with the intermediate filament cytoskeleton of the apical cap, which we exploited to increase the specificity of our BioID experiments (Figure S2B). We grew AC9^BioID^ parasites in biotin, lysed them in 1% Triton-X-100, and separated the cytoskeleton from solubilized membrane and cytosolic components by sedimentation at 14,000 × g. We then purified the biotinylated proteins using streptavidin resin, which were identified by LC-MS/MS (Supplemental Dataset S1). Our dataset was of high quality, as the top candidates were enriched in known apical cap proteins (12, 19). In addition, our top hit was previously undescribed (TGGT1_292950). To validate that TGGT1_292950 is, indeed, an apical cap protein, we epitope-tagged the endogenous gene with 3xHA. The resulting protein colocalizes with the apical cap marker ISP1 (Figure S2C), leading us to name the gene apical cap protein 10 (AC10).

Among the top candidates from our BioID dataset was the MAP kinase, ERK7 (TGGT1_233010). We recently demonstrated that ERK7 localizes to the apical cap, and that its loss-of-function phenotype is essentially identical to what we have observed for AC9 (17). We therefore prioritized investigating the function of this interaction. AC9 and ERK7 colocalize at the apical cap (Figure 3A) and proximity ligation immunofluorescence (34) demonstrated that they are closely associated at this site (Figure 3B). ERK7 is a member of an early-branching MAPK family that is conserved throughout eukaryotes (35, 36) and has been implicated as a facilitator of ciliogenesis in metazoa (15, 16) and Apicomplexa (17), though little is known about its signaling cascade and regulatory interactions in any organism. AC9 is conserved among coccidian parasites, though the overall protein sequences are highly divergent (21-55% identity). The region of highest conservation is in the AC9 C-terminus (Figure 3C). Analysis across all sequenced coccidian genera demonstrated that 16 of the most C-terminal 33 residues in AC9 are invariant.

**Figure 3:**
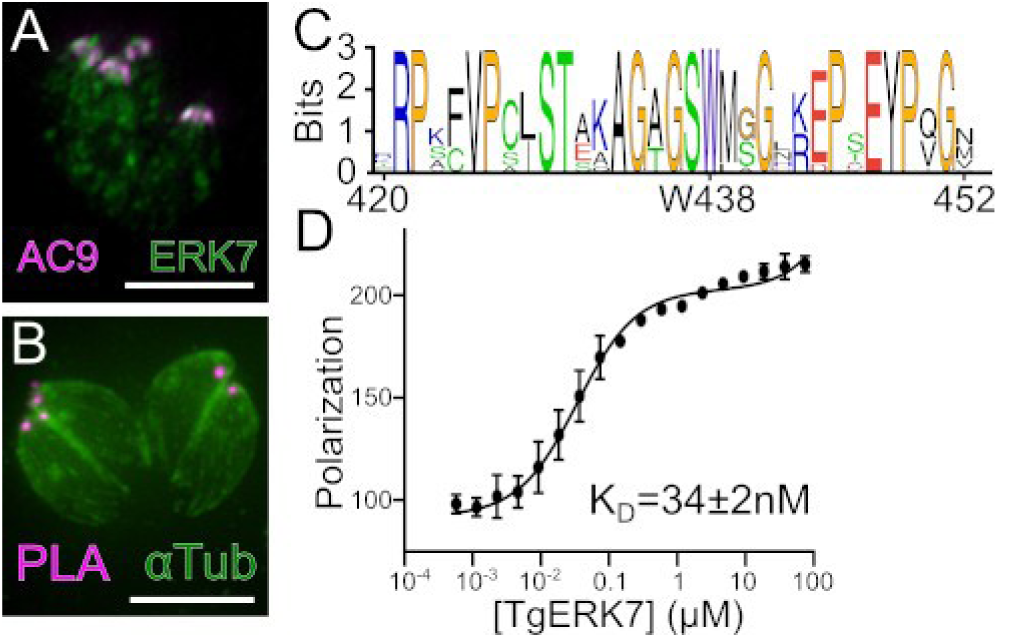
AC9 tightly binds ERK7. (A) ERK7-Ty (green) co-localizes with AC9-HA (magenta) at the apical cap. (B) Proximity ligation (magenta) of ERK7 and AC9 reveal bright foci at the parasite apical cap. Parasites are counterstained with anti-β-tubulin (Note that this antibody does not stain the apical complex, likely due to antigen accessibility). Scale bars: 5 μm. (C) Sequence logo for AC9_419-452_ highlights invariant C-terminal residues. (D) Binding of AC9_419-452_ to ERK7_1-358_ was measured by fluorescence polarization and the K_D_ was calculated from global fit of 3 replicate experiments of 3 technical replicates each.

To test whether the AC9 C-terminus was indeed the site of ERK7 binding, we bacterially expressed and purified several AC9 constructs N-terminally fused to yeast SUMO as a carrier protein. We found that AC9_401-452_ was robustly pulled-down by the GST-ERK7 kinase domain (Figure S3), demonstrating that the two proteins interact directly. The AC9 C-terminus is predicted to be intrinsically disordered, suggesting that it associates with the ERK7 kinase domain in an extended conformation. MAPKs interact with their substrates and regulatory partners through docking interactions that involve short linear motifs, usually 10-15 residues (37–39). We reasoned AC9 may be interacting through such a motif and tested whether shorter regions of the AC9 C-terminus were sufficient to bind ERK7. Surprisingly, we found that neither AC9_401-430_ nor AC9_431-452_ were detectable after pull down by GST-ERK7 (Figure S3). We therefore reasoned that the entire well-conserved portion of AC9, comprising AC9_419-452_ (Figure 3C), was required for the interaction. We measured the binding of fluorescein-labeled AC9_419-452_ to recombinantly expressed and purified ERK7 kinase domain by fluorescence anisotropy. AC9_419-452_ bound ERK7 with a K_D_ of 34±2 nM (Figure 3D), which is an affinity consistent with a strong MAPK-docking site interaction (37, 40).

### AC9 is required for ERK7 localization and function in parasites

Given their interaction, we reasoned that AC9 may be an ERK7 substrate. Available phosphoproteomic data report 9 phosphosites on AC9 (ToxoDBv46, http://toxodb.org; (41)). To test the relevance of these sites to our observed phenotypes, we complemented the AC9^AID^ strain by expressing a nonphosphorylatable allele, in which each of these Ser/Thr had been mutated to Ala (Figure S4A). This mutant protein correctly localized to the apical cap (Figure S4B) and fully rescued the ability of the AC9^AID^ parasites to form plaques (Figure S4C), demonstrating that phosphorylation on these sites is not required for AC9 function.

We identified ERK7 as a component of the cytoskeletal fraction of AC9-interacting proteins (Supplemental Dataset 1, Figure S2), but ERK7 has no domains that would be predicted to interact with cytoskeleton on its own. We therefore reasoned that AC9 may act as a scaffold that recruits ERK7 to the apical cap cytoskeleton. Consistent with this model, the ERK7 protein was lost from the apical cap upon AC9 degradation (Figure 4A), and this localization is rescued in the AC9 wt-complement (Figure 4B). Importantly, AID-mediated degradation of ERK7 has no effect on AC9 localization (Supplemental Figure S5A). These data suggest that a major function of AC9 is to recruit ERK7 to the apical cap. To directly test this model, we sought an AC9 mutant that retained its own localization but did not bind ERK7. While we were unable to obtain stable strains expressing deletions of the AC9 C-terminus, we found that mutation of 3 Ser/Thr to Glu (AC9^3xGlu^; Ser419Glu, Thr420Glu, Ser437Glu) reduced *in vitro* AC9 affinity for ERK7 by ∼200-fold (Figure 4C). AC9^3xGlu^ localized correctly when expressed in the background of the AC9^AID^ strain (Figure 4D,E), though it was unable to rescue the AC9^AID/IAA^ plaque phenotype (Figure 4F). In addition, while ERK7 localized to the apical cap in AC9^AID/3xGlu^ parasites, this localization was lost upon addition of IAA and degradation of the wild-type AC9^AID^ copy (Figure 4D), as was the tubulin-rich conoid (Figures 4E, S5B). Therefore, interaction with AC9 is necessary for ERK7 concentration at the apical cap. Importantly, ERK7 protein was still found in the parasite cytosol after AC9 degradation, demonstrating that expression of ERK7 is not sufficient for conoid development (Figures 4E, S5B). These data suggest that ERK7 must be present at the apical cap to function, and demonstrate that a critical function for AC9 is to control ERK7 localization.

**Figure 4.**
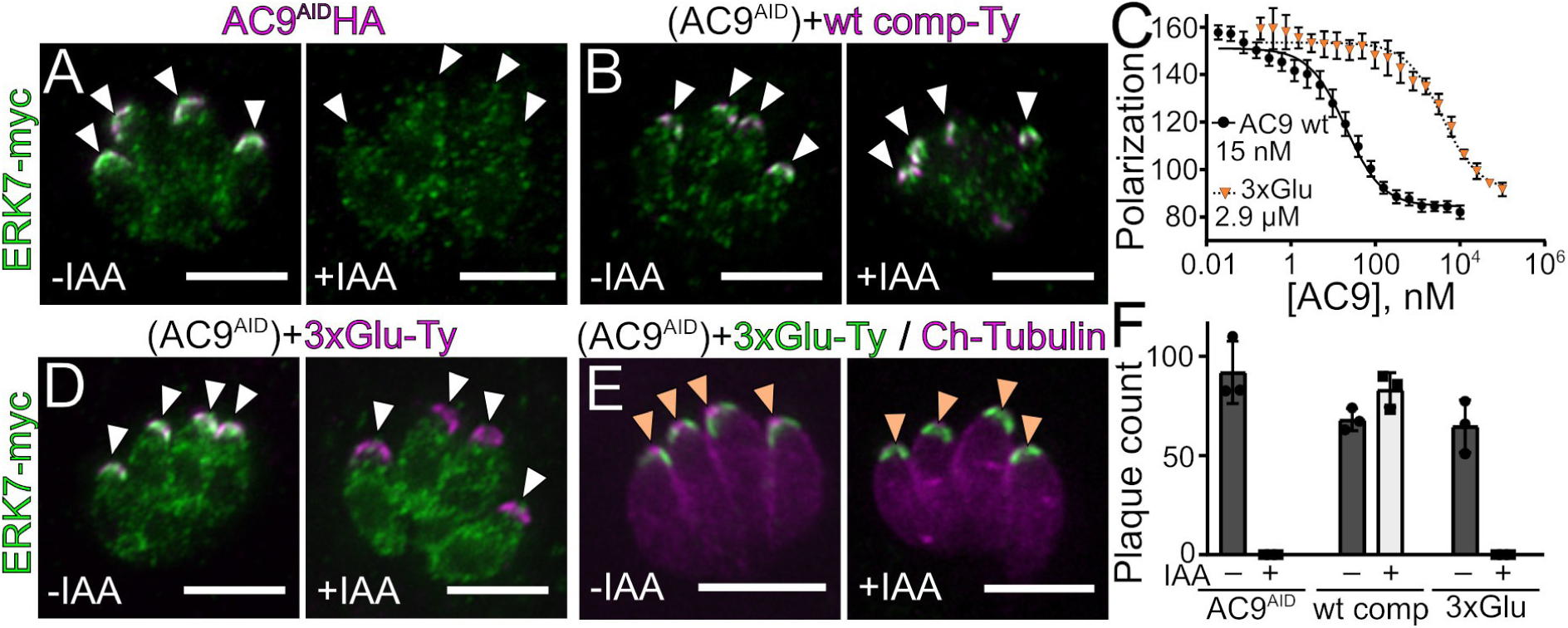
AC9 is a scaffold that drives ERK7 apical localization. (A) ERK7-3xmyc (green) localization is lost upon degradation of AC9^AID^-3xHA with growth in IAA (magenta), which is rescued in the (B) AC9 wt complement. White arrows indicate apical cap. (C) K_I_ of wt AC9_401-452_ and 3xGlu AC9_401-452_ were determined by competition with fluorescein-labeled AC9^419-452^. 95% CI wt: 13.4-17.7 nM; 3xGlu: 2.4-3.6 μM (D) ERK7-3xmyc (green) localization was compared in 3xGlu-complemented (magenta) parasites as in (A,B). (E) 3xGlu-complemented (green) AC9^AID^ parasites expressing mCherry-tubulin (magenta) were grown with or without IAA. Orange arrowheads indicate expected location of conoid foci. (G) AC9 localization is unaffected by ERK7 degradation. Scale bars: 5 μm.

### AC9 binds ERK7 in an unusual inhibitory conformation

To better define the nature of the AC9:ERK7 interaction, we solved the co-crystal structure of AC94_19-452_ bound to the ERK7 kinase domain to 2.1Å (Figure 5, Table S1). To our surprise, the AC9 C-terminus forms extensive contacts with ERK7 and wraps around the kinase from the MAPK docking site (the CD domain) to the substrate-recognition region (Figure 5). Strikingly, AC9-binding encompasses all major MAPK interacting and regulatory regions except for the F-site (42). MAPK docking site interactions are typically defined by a cluster of positively charged residues in the docking motif that interact with an acidic cluster in the kinase CD domain adjacent to Van der Waals interactions between complementary hydrophobic surfaces in the two partners (37, 43, 44). Remarkably, the only MAPK-interacting protein from *Toxoplasma* previously described is a secreted effector, GRA24, that binds mammalian p38 with a canonical docking site interaction (45); there are no regulatory partners known for the parasite’s MAPKs.

**Figure 5:**
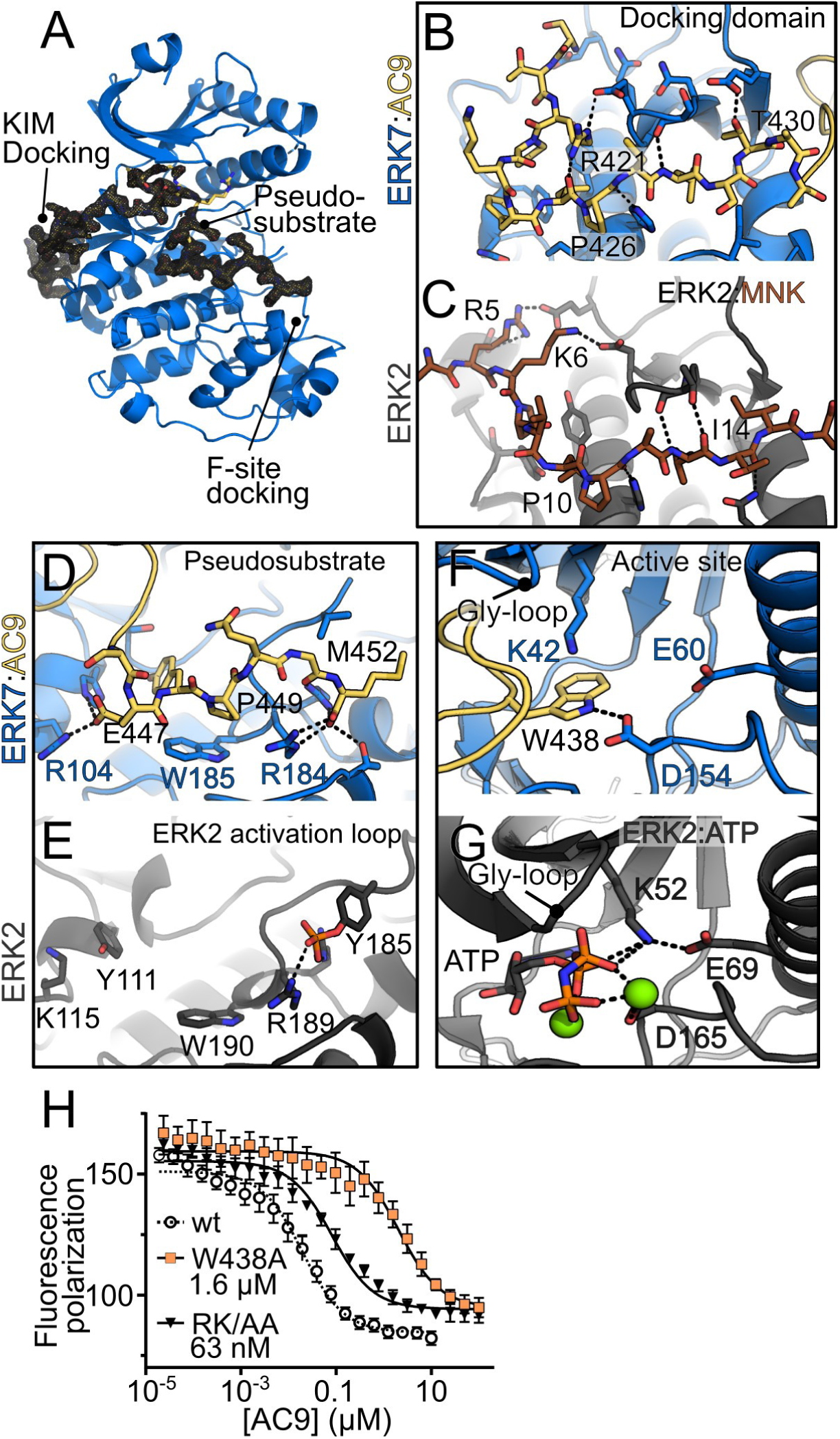
AC9 binds ERK7 in an inhibitory conformation. (A) Overview of AC9:ERK7 interaction. ERK7 is blue. A 1.5σ 2F_0_-F_C_ electron density map (black) is shown around AC9 (yellow). Contacts between AC9 (yellow) and ERK7 (blue) are compared with ERK2 (gray) at (B,C) the MAPK docking domain, (D,E) the activation loop/substrate binding site, and (F,G) the kinase active site. (H) K_I_ of AC9 mutants were determined by competition with fluorescein-labeled wild-type AC9. Wild-type competition curve (K_I_ = 15 nM) shown for comparison. 95% CI: AC9_W438A_ 1.3-2.0 μM; AC9_R421A/K423A_ (RK/AA) 55-74 nM. ERK2 images are from PDB 4H3Q (C) and 6OPG (E,G).

In stark contrast to more typical docking-site interactors, AC9 does not make extensive side-chain interactions with the MAPK CD domain. AC9 Phe424, Val425, Leu428 make weak van der Waals interactions with the ERK7 CD domain (contacts > 3.5Å). Only two polar AC9 side chains make close contact with the ERK7 CD domain (Figures 5B,S6): AC9 Arg421 is salt bridged with ERK7 Glu147 (3.0 Å), while AC9 Thr430 hydrogen bonds with ERK7 Glu96 (2.3 Å). Most of the ERK7 side-chains in the CD-domain that interact with AC9 do so through backbone hydrogen bonds. Mutation of the “basic patch” (*e.g.* MNK Arg5,Lys6 in Figure 5C) of typical motifs that bind the MAPK CD-domain abrogates their binding (37). Consistent with the idea that AC9 forms a sub-optimal ERK7 docking site interaction, mutation of the two basic residues in the AC9 CD-interacting motif (Arg421 and Lys423) to Ala only reduces affinity for ERK7 by only ∼4-fold (Figure 5H).

The C-terminal 8 residues of AC9 (444-452) appear to act as a pseudosubstrate, as these residues make close contacts with the ERK7 substrate recognition subdomain (Figure 5D). These interactions are centered around the invariant AC9 Pro449, which appears to mimic the required Pro in a true MAPK substrate and packs against ERK7 Trp185 (3.4 Å). Intriguingly, the C-terminus of AC9 makes contacts with a conserved basic cluster (ERK7 Arg184, 2.8 Å; Arg187, 2.9 Å) adjacent to the kinase APE motif. This cluster normally coordinates the pTyr in active MAPKs (Figure 5E), indicating AC9-binding partially displaces the ERK7 activation loop. Notably, the ERK7 binding motif in AC9 is of an invariant length at the C-terminus in all AC9 sequences (Figure 3C) and the interaction with the AC9 C-terminal carboxylate helps explain this conservation.

Most striking of the AC9 interactions with ERK7, however, is the insertion of AC9 Trp438 into the ERK7 active site (Figure 5F). The Trp438 side chain has displaced nucleotide and Mg^2+^ in our structure; even though the crystals formed in 1 mM ADP and 10 mM MgCl_2_, there is no density consistent with either. In fact, the Trp438 side-chain appears to be acting as a nucleotide mimic, a surprising function for a protein residue. The Trp438 indole ring makes π-cation interactions (3.2 Å) with the catalytic VAIK Lys42 that normally coordinates the ATP β-phosphate. In addition, the indole ring hydrogen bonds with the DFG Asp154 (2.7 Å) which normally coordinates nucleotide through a bound Mg^2+^. The AC9 loop flanking the Trp438 inserts from above the ERK7 CD domain, pushing the kinase Gly-loop into an open conformation without disrupting its overall secondary structure. Thus, the AC9-bound conformation of the ERK7 active site is substantially different from that typical of an active kinase, such as ERK2 (Figure 5G, S7A,B). Furthermore, our structure fully explains the loss of affinity of the AC9^3xGlu^ mutant for ERK7. While neither Ser419 nor Thr420 make side chain contacts, Ser437 is buried in the active site next to Trp438. Mutation of Ser437 to Glu, however, would clash with ERK7 Asp98 (Figure S7C), consistent with the 200-fold drop in affinity we observed.

Taken together, our structural data strongly suggested that AC9 acts not only as a scaffold for ERK7, but likely serves as a competitive inhibitor of its kinase activity. Indeed, we found that *Toxoplasma* ERK7 phosphorylation of the generic substrate myelin basic protein (MBP) was completely blocked by the addition 10 μM AC9_401-452_ to the reaction (Figure 6A). Because the interactions AC9 makes with *Toxoplasma* ERK7 side-chains are broadly conserved among MAPKs (Figure S6), we reasoned AC9 may be a promiscuous inhibitor. Surprisingly this was not the case. While AC9 robustly inhibits *Toxoplasma* ERK7 activity, it had no effect on the ability of another *Toxoplasma* MAPK (TgMAPK2), rat ERK7, or rat ERK2 to phosphorylate MBP (Figure 6A). It thus seems likely that the specificity of AC9 for TgERK7 is due to a combination of surface complementarity and the ability of TgERK7 to adopt a conformation able to recognize AC9.

**Figure 6:**
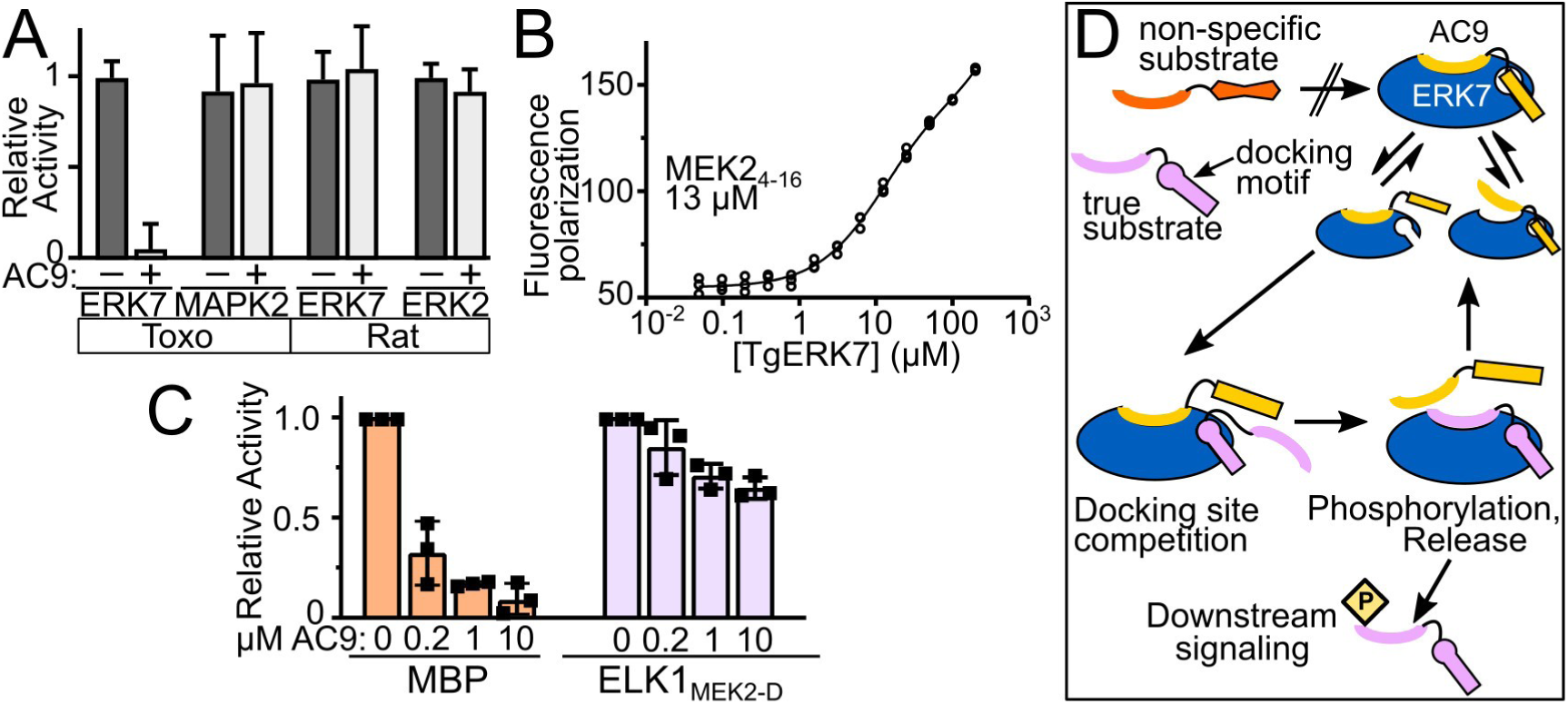
AC9 is an inhibitory regulator of ERK7 kinase activity. (A) Quantification of phosphorylation of MBP by the indicated kinases in the presence and absence of 10 μM AC9_401-452_, normalized to activity without AC9 (n=3). (B) Binding of the KIM from rat MEK2 to TgERK7 was measured by fluorescence polarization. 95% CI: 10.5-14.5 μM. (C) Quantification of TgERK7 phosphorylation of MBP or a chimeric ELK1/MEK2_4-16_ substrate in the presence of the indicated AC9_401-452_ concentration normalized to activity without AC9 (n=3). (D) Model for AC9 regulation of ERK7 specificity. AC9 occupies the ERK7 active site, preventing the binding of non-specific substrates. However, the AC9 docking-site interaction is sub-optimal and can be competed off by true ERK7 substrates, which are released after phosphorylation, allowing AC9 to rebind. All error bars are S.D.

As described above, the AC9 binds ERK7 with an extended interface that buries ∼1650 Å^2^. While the AC9:ERK7 interaction is reasonably strong (15-35 nM; Figures 3D, 4C), protein:peptide interactions can achieve similar affinities with substantially smaller buried surfaces. For instance, Grb2-SH2 binds Shc1_423-435_ with a K_D_ of 18 nM (46) with a 404 Å^2^ interface (47). Similarly, the Cbl-TKB domain binds APS_609-621_ with a K_D_ of 43 nM with a 629 Å^2^ interface (48). These data suggest the AC9:ERK7 interface is formed by a distributed series of weak interactions, which is consistent with our mutagenesis data. Mutation of residues usually critical to recognizing the MAPK CD-domain only modestly affected AC9 affinity (AC9_R231A/K232A_; Figure 5H). Even mutation of Trp438 yields a protein with a respectable, albeit weakened, affinity of 1.6 μM (Figure 5H). In addition, these distributed interactions (CD-domain, active site, activation loop/pseudosubstrate) appear to act cooperatively to enable AC9 to capture ERK7 in an optimal conformation for binding. Truncated versions of AC9 showed no measurable binding to the kinase (Figure S3).

Given these observations, we reasoned that AC9 may be displaced by a substrate that can engage the CD-domain with a physiologically relevant affinity. While there are yet no known substrates of *Toxoplasma* ERK7, we tested available peptides with known affinities to metazoan MAPKs for ERK7-binding. Even though there is no homologous sequence in *Toxoplasma*, we found that rat MEK2_4-16_ binds *Toxoplasma* ERK7 with a K_D_ of ∼12 μM (Figure 6B). Note that physiological MAPK docking site interactions range in affinity from 0.5 – 50 μM (40, 49, 50) and MEK2 engages its cognate partner, ERK2, with an ∼8 μM K_D_. We chose to modify a well-characterized mammalian MAPK substrate, ELK1, which requires CD-domain interaction for efficient phosphorylation (51). As expected, we could not detect measurable phosphorylation of an ELK1_300-428_ construct, which lacks an ERK7-binding motif. A chimeric ELK1 construct that contains the MEK2 docking motif (ELK1_MEK2-D_), however, was phosphorylated by ERK7 at levels similar to the generic substrate MBP in the absence of AC9 (Figure 6C). While ERK7 phosphorylation of MBP was efficiently inhibited by AC9, substantial levels of ELK1_MEK2-D_ phosphorylation remained even in the presence of 10 μM AC9 (Figure 6C). Thus AC9 inhibition of ERK7 can be successfully released by competition with even a modest-affinity docking-site interaction.

Notably, ERK7 family members are unusual in the MAPK family as they are able to autophosphorylate their activation loops (18), and thus bypass the need for a MAPK kinase for their activation. We have previously found that ERK7 kinase activity is required for conoid formation (17). Our data therefore suggest that AC9 has the dual roles of concentrating *Toxoplasma* ERK7 at the apical cap and regulating its kinase activity and substrate specificity (Figure 6D).

## Discussion

We have identified AC9 as an essential cytoskeletal scaffold of the *Toxoplasma* ERK7 kinase. Remarkably, inducing AC9 degradation phenocopies loss of ERK7 and results in parasites that mature without conoids, thereby interrupting the lytic cycle. We found that AC9 is required for ERK7 localization to the apical cap and identified a mutant that could not bind and recruit ERK7, and therefore did not rescue any of the AC9 loss-of-function phenotypes (Figure 4). Surprisingly, our crystal structure of the ERK7:AC9 complex revealed that AC9 is also a competitive inhibitor of ERK7 kinase activity. Usually, genetic ablation of an inhibitor would be expected to present the opposite phenotype from its target. Because ERK7 kinase activity is required for conoid formation (17), our data suggest that AC9 inhibition of the kinase is not permanent. Instead, we propose that AC9 binding to ERK7 represents an unusual mechanism of ensuring kinase specificity (Figure 6D). Because ERK7 is autoactivating, its regulation by a phosphatase would be insufficient to ensure signaling fidelity, especially when the kinase is maintained at a high local concentration at the apical cap. While the full AC9 C-terminus binds tightly to ERK7 (Figure 3D), this is due to cooperative binding of distributed contacts (Figure 5); neither AC9^401-430^, which occupies the CD-domain, nor AC9^431-452^, which occupies the active and pseudosubstrate sites, are sufficient for ERK7 binding (Figure S3). We therefore propose that an ERK7 substrate with both a strong kinase-interacting motif and substrate site would successfully compete with AC9 for kinase binding during phosphorylation (Figure 6D). Alternatively, an ERK7-activating factor may transiently displace AC9, allowing substrate binding. In either of these cases, the regulated release of AC9 inhibition would define ERK7 activity. Determining ERK7 substrates as well as other regulatory factors will not only provide the opportunity to directly test these models, but will also allow us to define how ERK7 facilitates ciliogenesis in parasites (17) and other organisms (15, 16).

A striking feature of the AC9 inhibition of ERK7 is its specificity. Even though AC9 makes contacts with conserved sites, we found that it does not inhibit other *Toxoplasma* or mammalian MAPKs, including the rat ERK7 ortholog. These data suggest that the kinase dynamics, rather than final contact sites, are contributing to specificity of the inhibition. This has clear implications for the design of specific inhibitors to parasite ERK7. Moreover, the AC9:ERK7 interaction may represent a generalized mechanism of inhibiting ancient signaling molecules such as ERK7. ERK7 is the earliest branching member of the MAPK family (52), and is unusual in its auto-activation (18). Notably, cyclin-dependent kinases (CDK) are also regulated by inhibitory proteins (53), and are related to the MAPKs (52). Like the ERK7:AC9 interaction, the cell cycle inhibitor p27 is intrinsically disordered and binds CDK2 through a distributed surface. Also like AC9, p27 inserts itself into the kinase active site and displaces nucleotide (Figure S7C) (54). However, there are notable differences between the two inhibitory interactions. First, AC9 occupies the MAPK CD-domain, which is not present in CDKs. Also, while our structure demonstrates that AC9 insertion pushes the ERK7 Gly-loop into an inactive conformation (Figures 5,S7), p27 replaces the first β-strand in CDK2, destroying the Gly-loop structure (Figure S7C). Finally, while both ERK7 and AC9 protein levels appear to be maintained stably at the apical cap throughout the cell cycle, p27 is degraded at the G_1_/S transition, releasing its inhibition of CDK2 (55). Nevertheless, the convergent evolution of specific protein inhibitors of two branches of the CMGC family suggests that there may be additional genetically encoded kinase inhibitors that remain unidentified throughout Eukaryota. Identifying such inhibitors would refine our understanding of cellular signaling architecture and provide potential platforms on which specific therapeutics may be designed.

## Materials and Methods

### Sequence analysis

Sequences for AC9 from *Toxoplasma gondii, Hammondia hammondi, Neospora caninum, Cyclospora cayetanensis, Cystoisospora suis, Eimeria spp.*, and *Sarcocystis neurona* were retrieved from ToxoDBv43 (https://www.toxodb.org) using BLAST. The AC9 sequence from *Besnoitia besnoiti* was identified by BLAST and retrieved from Uniprot. The protein sequences were aligned with Clustal Omega. (56). The AC9 sequence logo was generated with WebLogo (57).

### PCR and plasmid generation

All PCR was conducted using Phusion polymerase (NEB) using primers listed in Supplemental Dataset 2. Constructs were assembled using Gibson master mix (NEB). Point mutations were created by the Phusion mutagenesis protocol. AC9^AID^ tagging constructs were generated using a PCR-amplified homology-directed repair template (P1-2) and a CRISPR/Cas9 pU6-Universal plasmid with a protospacer against the 3’ UTR of the gene of the interest (P3-4). AC9-BirA*, ERK7 and AC10 C-terminal tagging constructs were generated using P1-4, P5-8 and P9-12, respectively. To generate the wildtype complementation construct, the complete AC9 coding sequence was PCR-amplified from cDNA and cloned into a UPRT-locus knockout vector driven by the ISC6 promoter. Both the 3xGlu and the phosphorylation mutants were constructed using synthetic genes (Quintara Biosciences) and cloned into the pISC6-UPRT vector (Supplemental Dataset S2).

### Chemicals and Antibodies

3-indoleaceticacid (IAA; heteroauxin; Sigma-Aldrich I2886) was used at 500 µM from a 500 mM stock in 100% ethanol. A23187 (Sigma-Aldrich C5722) was used at 1-2 μM, dissolved from a 2 mM stock dissolved in DMSO. The HA epitope was detected with mouse monoclonal antibody (mAb) HA.11 (BioLegend MMS-101P, San Diego, CA), rabbit polyclonal antibody (pAb) anti-HA (Invitrogen 71-5500), or rat monoclonal 3F10 (Sigma-Aldrich 11867423001). The Ty1 epitope was detected with mouse mAb BB2 (58). The c-Myc epitope was detected with mouse mAb 9E10 (59) or rabbit pAb anti-Myc (Invitrogen PA1981). His_6_-tagged proteins were recognized with mouse anti-His_6_ (R&D systems MAB050). Biotinylated proteins were detected with Alexa-fluor-488-streptavidin (Molecular Probes S32354) and captured with Streptavidin High-Capacity Agarose beads (Thermo Scientific PI20359) *Toxoplasma*-specific antibodies include rabbit anti-β-tubulin (17), mouse mAb anti-ISP1 7E8 (11), rabbit pAb anti-SAG1 (60), mouse mAb anti-F1-ATPase beta subunit 5F4 (61), mouse mAb anti-MIC2 (gift from Vern Carruthers), rat pAb anti-GRA39 (62), rabbit pAb anti-ROP2, rat pAb anti-RON11 (61), mouse mAb anti-SAS6L (4), mouse mAb anti-IMC1 45.15 (63), and mouse mAb anti-ISP3 (11).

### Immunofluorescence

HFF cells were grown on coverslips in 24-well plates until confluent and were infected with parasites. The cells were rinsed once with phosphate buffered saline (PBS), fixed with 3.7% formaldehyde in PBS, washed, permeabilized with 0.1% Triton-X-100, blocked with 3% BSA for 1 hour, and incubated with primary antibodies for a minimum of 2 hours. Secondary antibodies used were conjugated to AlexaFluor-488 or AlexaFluor-594 (ThermoFisher). The coverslips were mounted in Vectashield (Vector Labs) and viewed with an Axio Imager.Z1 fluorescent microscope (Zeiss). For proximity ligation assays, cells were fixed in 4% paraformaldehyde / 4% sucrose followed by permeabilization in 0.1% Triton-X-100 for 10 min. Blocking, antibody incubations and proximity ligation were conducted according to manufacturer’s directions (Sigma-Aldrich) using rabbit polyclonal anti-HA (64) and mouse m2 anti-FLAG (Sigma-Aldrich F1804) as primary antibodies. Cells were counterstained with AlexafFluor-647 conjugated rabbit anti-β-tubulin, mounted in VectaShield, and imaged on a Nikon Ti2E microscope.

### Western blotting

Parasites were lysed in Laemmli sample buffer with 100 mM DTT and heated at 100°C for 10 min. Proteins were separated by SDS-PAGE, transferred to nitrocellulose, and probed with primary antibodies and the corresponding secondary antibody conjugated to horseradish peroxidase. Western blots were imaged using the SuperSignal West Pico substrate (Pierce) and imaged on a ChemiDoc XRS+ (Bio-Rad, Hercules, CA). Band intensities were quantified using the manual volume tool in the Image Lab Software of the ChemiDoc XRS+.

### Plaque assays

HFF monolayers were supplemented with -/+ 500 µM IAA before allowing equal numbers of freshly lysed, extracellular parasites of a given strain (grown in -IAA) to infect and form plaques for 7 days. Cells were then fixed with ice-cold methanol and stained with crystal violet. Plaque number was counted manually and analyzed by Prism GraphPad. All plaque assays were performed in triplicate for each condition.

### Invasion assays

Invasion assays were performed as previously described (22). Parasites were grown for 30 hours -/+ IAA and intracellular parasites were collected by scraping and passaging through a 27-gauge needle. Equivalent parasite numbers were resuspended in Endo buffer (65) and settled onto coverslips with HFF monolayers for 20 minutes. Endo buffer was then replaced with warm D1 media and incubated at 37°C for 30 minutes. Coverslips were then fixed, blocked, and extracellular parasites were stained with anti-SAG1 antibodies. The samples were then permeabilized, and all parasites stained with anti-F1B ATPase antibodies, and incubated with secondary antibodies. Parasites were scored as invaded (SAG1+, F1B-ATPase-) or not (SAG1+, F1B+) by fluorescence microscopy. Invasion assays were performed in triplicates, at least 10 fields were counted for each replicate, and the average for each replicate was calculated as a percentage.

### Egress assays

Parasites were grown on a monolayer on coverslips for 34 hours -/+ IAA until most vacuoles contained 16 or 32 parasites. Coverslips were washed twice with pre-warmed PBS and incubated with A23187 (or DMSO control) diluted in PBS at 37°C for 2 minutes. Coverslips were then fixed and stained with rabbit anti-IMC12 antibodies (P. Back, in preparation). At least 10 fields of ∼200 vacuoles/field were counted for 3 replicate coverslips for each condition.

### Microneme secretion

Microneme secretion assays were performed as previously described (23). Briefly, parasites were grown for 30 hours and intracellular parasites were collected by mechanical release through a 27-gauge needle. After washing twice in D1 media (DMEM, 20 mM HEPES, 1% FBS), parasites were resuspended in pre-warmed D1 media and induced with 1 µM A23187 for 10 minutes at 37°C. Secretion was arrested by cooling on ice, and parasites were pelleted 1000 x g for 5 minutes at 4°C. The supernatant was collected and centrifuged again 1000 x g. Secreted proteins in the resulting supernatant were assessed by SDS-PAGE and Western blot analyses.

### Detergent fractionation and streptavidin purification

The IMC cytoskeletal fraction was isolated by 1% Triton X-100 fractionation as described (66). Extracellular parasites were lysed in 1% Triton X-100, 50 mM Tris-HCl, pH 7.4, 150mM NaCl buffer supplemented with Complete Protease Inhibitor Cocktail (Roche) and incubated on ice for 30 min. Lysates were centrifuged, and equivalent loads of the total, supernatant, and pellet samples were run on SDS-PAGE and immunoblotted, using IMC1 and ISP3 as cytoskeletal and soluble controls, respectively. For streptavidin purification of BioID samples, parasites were grown for 24 h in media supplemented with 150 μM biotin. The cytoskeletal fraction was solubilized in 1% SDS and diluted to RIPA conditions, biotinylated proteins were purified using Streptavidin High-Capacity Agarose (Pierce), and the proteins were identified via mass spectrometry.

### Mass spectrometry

Purified proteins bound to streptavidin beads were reduced, alkylated and digested by sequential addition of lys-C and trypsin proteases (67, 68). Samples were then desalted using C18 tips (Pierce) and fractionated online using a 75 μM inner diameter fritted fused silica capillary column with a 5 μM pulled electrospray tip and packed in-house with 15 cm of Luna C18(2) 3 μM reversed phase particles. The gradient was delivered by an easy-nLC 1000 ultra high-pressure liquid chromatography (UHPLC) system (Thermo Scientific). MS/MS spectra was collected on a Q-Exactive mass spectrometer (Thermo Scientific) (69, 70). Data analysis was performed using the ProLuCID and DTASelect2 implemented in the Integrated Proteomics Pipeline-IP2 (Integrated Proteomics Applications, Inc., San Diego, CA) (71–73). Database searching was performed using a FASTA protein database containing *T. gondii* GT1 translated ORFs downloaded from ToxoDB on February 23, 2016. Protein and peptide identifications were filtered using DTASelect and required minimum of two unique peptides per protein and a peptide-level false positive rate of less than 5% as estimated by a decoy database strategy. Candidates were ranked by normalized spectral abundance factor (NSAF) values comparing the AC9^BioID^ versus control samples (74).

### Transmission electron microscopy

To prepare parasite ghosts for TEM, parasites were first incubated with 20 μM calcium ionophore in Hanks Buffered Saline Solution at 37°C for 10 min. The parasite suspension was allowed to adhere to a grid, after which membranes were extracted by addition of 0.5% Triton-X-100 for 3-4 min. The samples were then stained with 2% phosphotungstic acid, pH 7.4. All TEM images were acquired on a Tecnai G2 spirit transmission electron microscope (FEI) equipped with a LaB_6_ source at 120 kV.

### Protein expression and purification

Unless otherwise noted, proteins were expressed as N-terminal fusions to His_6_-SUMO. All proteins were expressed in Rosetta2 (DE3) bacteria overnight at 16°C overnight after induction with 300 mM IPTG. TgERK7 for crystallography and binding was co-expressed with λ-phosphatase. For His_6_-tagged proteins, cells were resuspended in 50 mM Tris, pH 8.6, 500 mM NaCl, 15 mM Imidazole and lysed by sonication. His_6_-tagged protein was affinity purified using NiNTA resin (Qiagen), which was washed with binding buffer. Protein was eluted in 20 mM Tris, pH 8.6, 100 mM NaCl, 150 mM Imidazole. ERK7_2-358_ and AC9 constructs were further purified as follows. Protein was diluted 1:1 with 20 mM Tris, pH 8.6 and purified by anion exchange on a HiTrapQ column. The resulting peaks were pooled, incubated with ULP1 protease for 30 min, after which they were diluted 1:1 in water and the cleaved SUMO separated from the protein of interest by anion exchange. The flow-through was concentrated and purified by size-exclusion chromatography, where it was flash frozen in 10 mM HEPES, pH 7.0, 300 mM NaCl for storage. GST and GST-ERK7 kinase domain (residues 2-358) affinity purified using glutathione sepharose (GE) and eluted with 10 mM glutathione, which was removed by dialysis in storage buffer before concentration and flash freezing.

### GST pulldowns

Purified GST and GST*-*ERK7_2-358_ were bound to glutathione sepharose (GE) in 10 mM HEPES, pH 7.0, 300 mM NaCl, 10 mM DTT. Equimolar amounts of purified SUMO fusions of AC9_401-452_, AC9_401-430_, or AC9_431-452_ (with an additional disordered linker composed of AC9_401-405_) were incubated with the glutathione resin for 5-10 min and washed 4× in binding buffer. The bound protein was removed by boiling in 1× SDS sample buffer, separated by SDS-PAGE, and detected by western blot analyses with mouse anti-His_6_ (Sigma).

### Fluorescence polarization

AC9_419-452_ with an additional N-terminal Cys was purified, as above, and allowed to react overnight with fluorescein-5-maleimide (Molecular probes). Free fluorescein was removed by sequential buffer exchange in an ND-10 desalting column and then in a 3 kDa MWCO centrifugal concentrator. Binding affinity was measured by serially diluting ERK7_2-358_ against 10 nM fluorescein-AC9 in 20 μL volumes in a 384 plate, and fluorescence polarization was measured in a BioTek Synergy plate reader. MEK2 binding was conducted in the same manner by titrating ERK7 against 100 nM fluorescein-labeled synthetic MEK2_4-16_ peptide (gift from Melanie Cobb). Competition experiments were conducted by titrating unlabeled AC9 constructs against 10 nM fluorescein-AC9 and 100 nM ERK7. All curves were globally fit in GraphPad Prism (to single-site binding or single-site inhibition, as appropriate) from biological triplicates of independent experiments composed of triplicate samples. 95% confidence intervals for all fit affinities are indicated in the legends.

### Protein crystallization

A 1:1 ERK7_2-358_:AC9_419-452_ complex at 9 mg/mL total protein with 10 mM MgCl_2_, 1 mM ADP, 10 mM DTT was mixed 1:1 in a sitting drop with 0.15 M DL-Malic acid, pH 7.0, 20% PEG 3350. Crystals were flash frozen in a cryoprotectant of reservoir with 25% ethylene glycol.

### Data collection, structure determination, and refinement

The diffraction data were collected at the UT Southwestern Structural Biology core with the Rigaku MicroMax-003 high brilliancy CuK-α X-ray generator, equipped with a Rigaku HyPix direct photon detector, and processed using the CrysAlisPro software package. A model of *Toxoplasma* ERK7 was created in Modeller v9.14 (75) using PDB 3OZ6 and used as a search model for molecular replacement in Phaser (76). Cycles of manual rebuilding in Coot (77) and refinement in Phenix (78), led to a final 2.1 Å structure of the ERK7:AC9 complex (PDB accession: 6V6A). The structure was evaluated with Molprobity (79).

### In vitro kinase assays

The kinase assays comparing specificity of AC9 inhibition were run using 1 µM of the indicated kinases, 5 mM MgCl_2_, 200 µM cold ATP, 10 mM DTT, 1 mg/mL BSA, 300 mM NaCl, 20 mM HEPES pH 7.0, 10% glycerol. TgERK7, rat ERK7, and TgMAPK2 were bacterially-expressed as His_6_-SUMO fusions and purified without phophatase treatment. The coding sequence for rat ERK7 was a gift of Marsha Rosner. Activated rat ERK2 was a gift of Melanie Cobb. Reactions were started by adding a hot ATP mix that contained 10 µCi γ[^32^P] ATP and 5 µg MBP. The 25 µL reactions were incubated at a 30°C water bath for 30 min. Reactions were stopped by adding 9 µL 4x SDS-buffer. 20 µL samples were then run on an SDS-PAGE gel. The gels were coomassie stained, and the MBP band was excised and radioactivity quantified using a scintillation counter. Recombinant ELK1 and ELK1_MEK2-D_ were expressed as His_6_-SUMO fusions and purified according to the same protocol as AC9. Competition assays were performed as above, with 200 nM TgERK7, 100 μM cold ATP, 10 μM of either MBP or ELK1_MEK2_ substrates, and varying concentrations of AC9_401-452_. These assays were imaged by phosphorimager (FujiFilm FLA-5100) and quantified using the ImageJ gel quantification tool (80).

### Figure generation

Data plotting and statistical analyses were conducted in GraphPad Prism v8.3. All error bars are mean-centered SD. All figures were created in Inkscape v0.92.

## Acknowledgments

We thank Betsy Goldsmith and Melanie Cobb for guiding discussions and Vasant Muralidharan, Vinnie Taggliabracci, and Ben Weaver for helpful comments on the manuscript; the UT Southwestern Electron Microscopy core facility for assistance with data collection and Zhe Chen and the UT Southwestern Structural Biology Lab for assistance with data collection and processing. X-ray data were collected on shared equipment funded by NIH grant S10 OD025018. M.L.R. acknowledges funding from the Welch Foundation (I-1936-20170325) and National Science Foundation (MCB1553334). X.H. was funded, in part, by Cancer Prevention and Research Institute of Texas Training Grant RP160157. P.J.B. acknowledges funding from the National Institutes of Health (NIAID R01 AI123360) and J.A.W. acknowledges NIH R01 GM089778. P.S.B. was funded by the Ruth L. Kirschstein NRSA GM007185.

## Supplemental Materials

**Figure S1.**
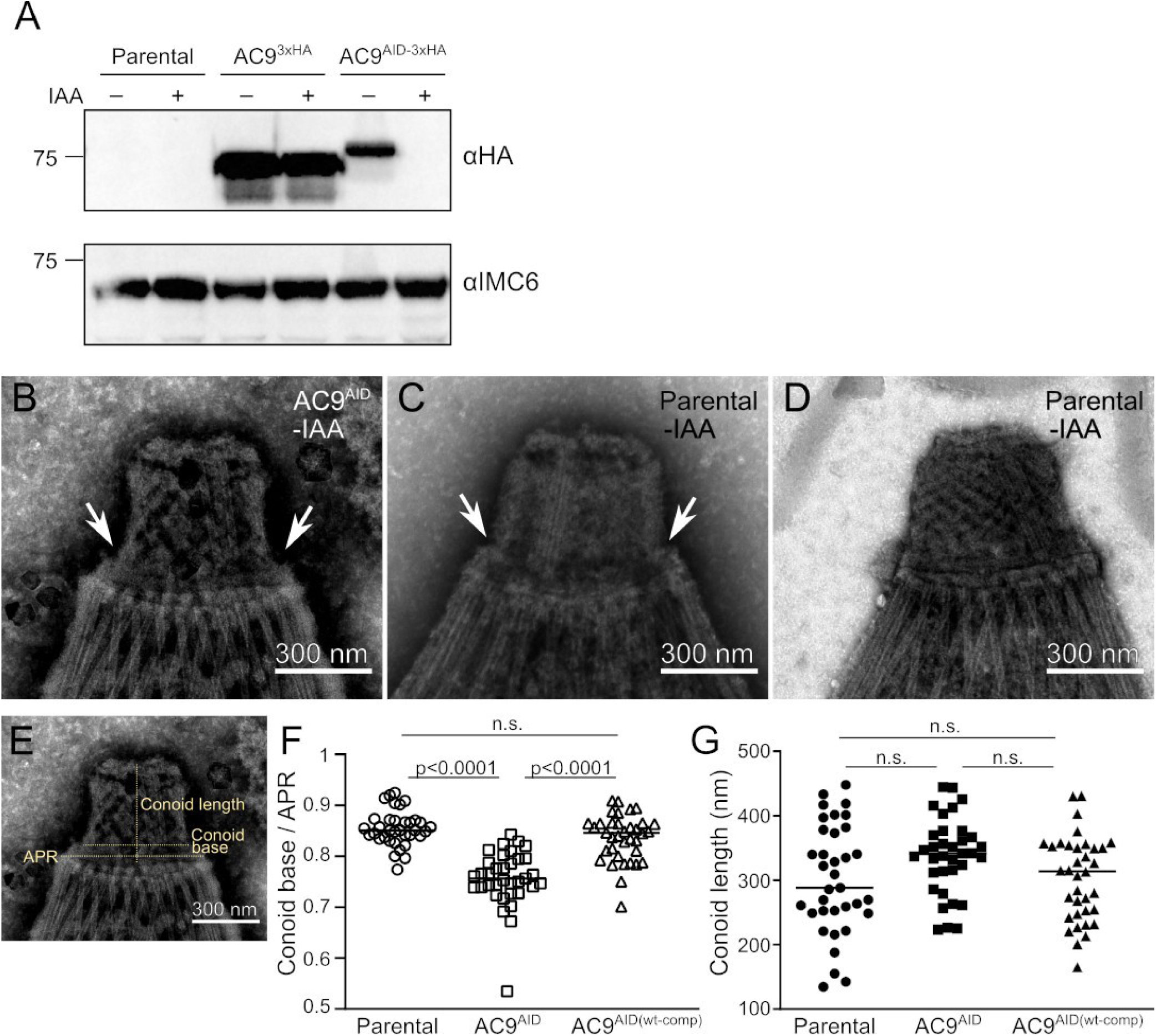
(A) IAA-independent reduction in AC9 protein levels upon AID tagging. Lysates from equal numbers of extracellular parasites of the indicated strains were separated by SDS-PAGE and probed with either anti-HA or anti-IMC6 (loading control). Quantification of band intensities indicates AC9^AID-3xHA^ levels are ∼39% of AC9^3xHA^ even when grown without IAA. Western blot also verifies undetectable levels of AC9 after overnight grown in IAA. (B) AC9^AID^ tagging (in the absence of IAA) exacerbates an ultrastructural artifact in TEM sample preparation (arrows) where conoid base slightly detaches from apical polar ring (APR). Note that similar preparations of parental parasites exhibit a similar phenotype (C), though most show stable conoids (D). This observation led us to quantify n=35 conoid TEMs from 2 independent sample preparations for each strain using the rubric in (E); note image is identical to (B). (F) There is a slight, but significant reduction in the ratio of the width of the conoid base over the APR width in AC^AID^ parasites as compared with parental and rescue strains. (G) We observed no significant difference in conoid extension lengths between the three strains. Significance estimated by 1-way ANOVA with Tukey’s test.

**Figure S2.**
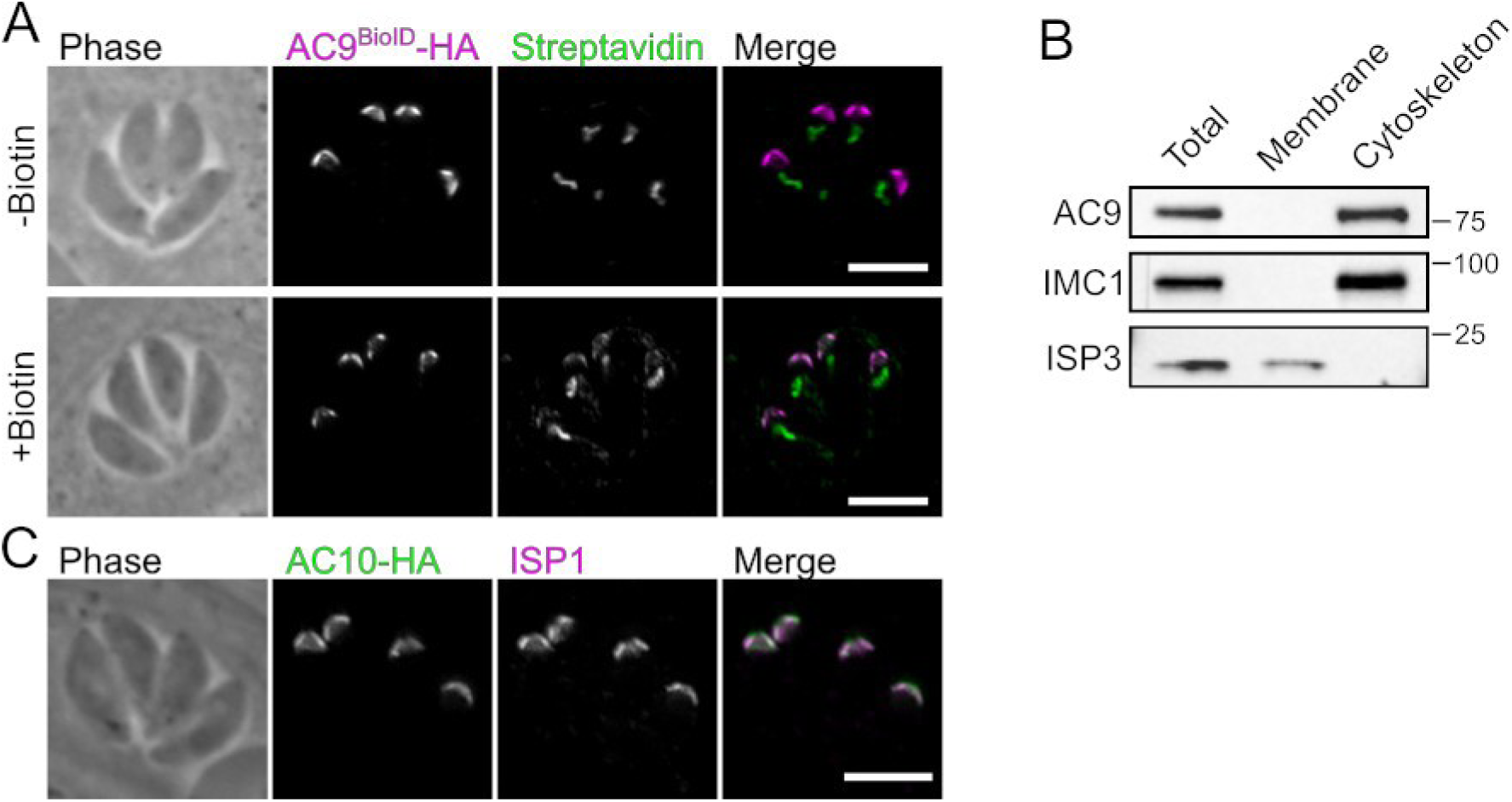
Preparation of apical cap cytoskeleton for BioID. (A) AC9^BioID^-3xHA (magenta) actively biotinylates proteins at the apical cap when parasites are grown in the presence of 150 μM biotin, as detected by streptavidin staining (green). Note that the parasite apicoplast contains natively biotinylated proteins that are recognized by streptavidin. (B) Cytoskeletal components of the apical cap such as AC9 and IMC1 are enriched by detergent fractionation, while membrane-associated apical cap proteins, such as ISP3, are de-enriched. (C) Of the top hits in our BioID dataset (Supplemental Data 1) was one previously unidentified apical cap protein, TGGT1_292950, which, when endogenously tagged at its C-terminus with 3xHA (green) was verified to localize to the apical cap (ISP1; magenta). All scale bars: 5 μm.

**Figure S3.**
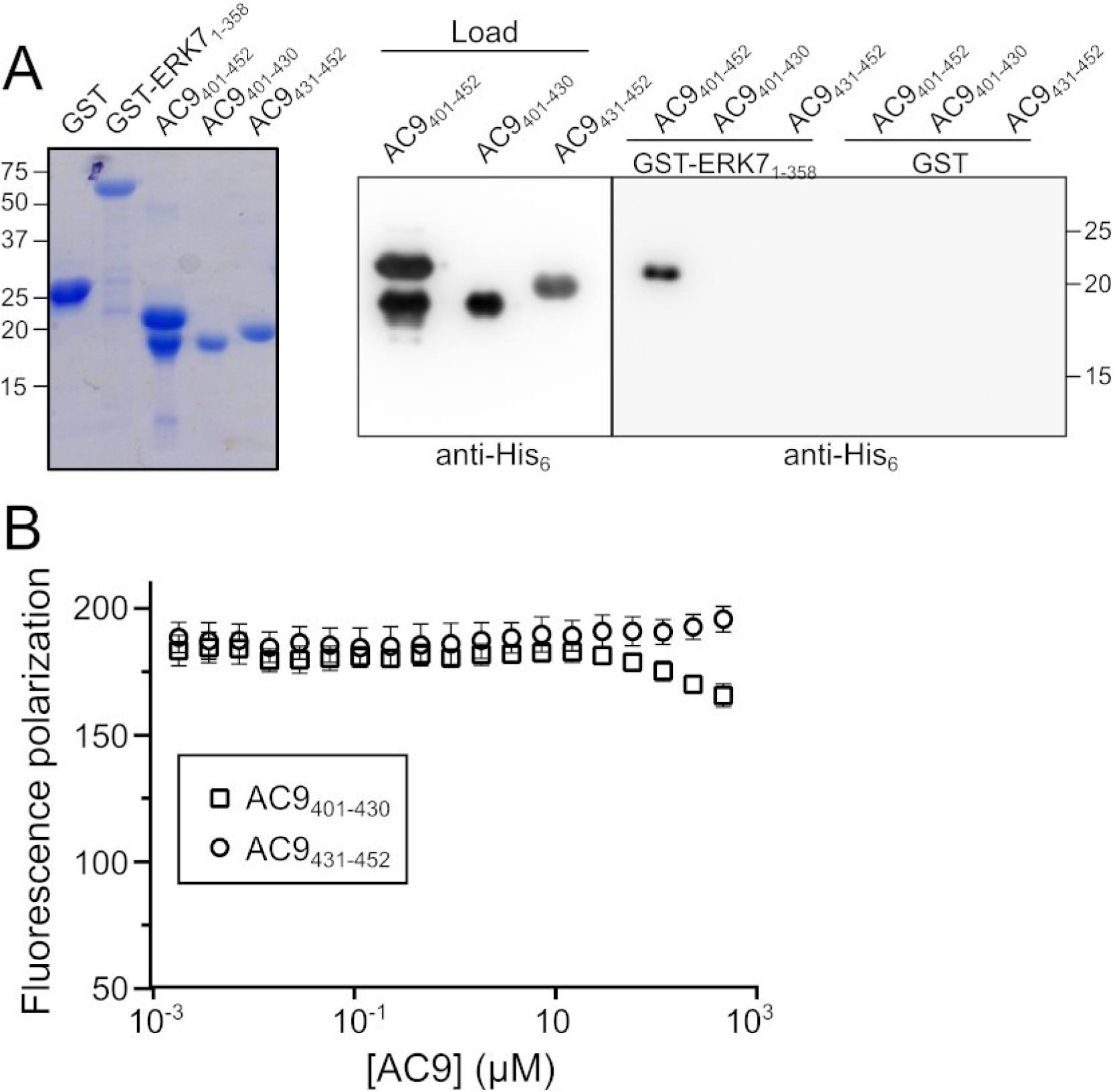
AC9 binds the ERK7 kinase domain. (A) (Left panel) Coomassie stained gel of purified proteins used to test binding. (Right panel) His_6_-SUMO fusions of the indicated AC9 fragments were incubated with GST or GST-ERK7 bound to glutathione sepharose resin, washed, and detected by western blot with anti-His_6_ antibody. Only AC9_401-452_ showed detectable binding to ERK7. (B) Fluorescence polarization competition in which AC9_401-430_ and AC9_431-452_ were titrated against fluorescein-labeled AC9_419-452_ bound to TgERK7. Note that neither the N-terminal nor C-terminal fragments of the AC9 ERK7-binding region were able to efficiently compete with AC9_419-452_ for binding, indicating the fragments have affinities >1 mM for the kinase.

**Figure S4.**
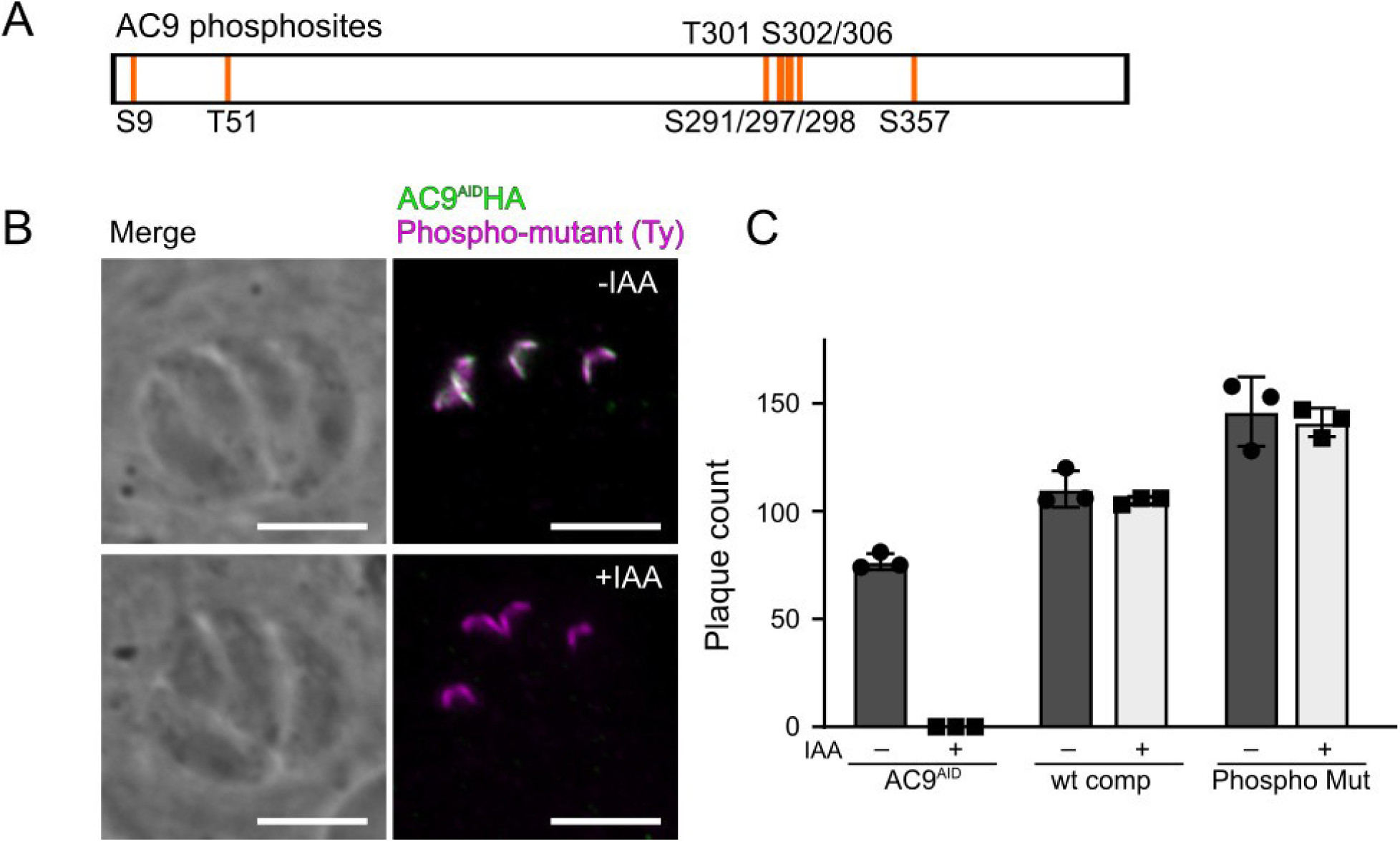
Phosphorylation of AC9 is not required for its function. (A) Location of known phosphorylated Ser/Thr on AC9 that have been mutated to Ala in this study. (B) Phospho-mutant AC9 correctly localizes to the apical cap in the AC9^AID^ background. Scale bars: 5 μm. (C) The phospho-mutant is able to fully rescue plaque formation in the AC9^AID^ background, indistinguishably from complementation with a wild-type copy of AC9.

**Figure S5.**
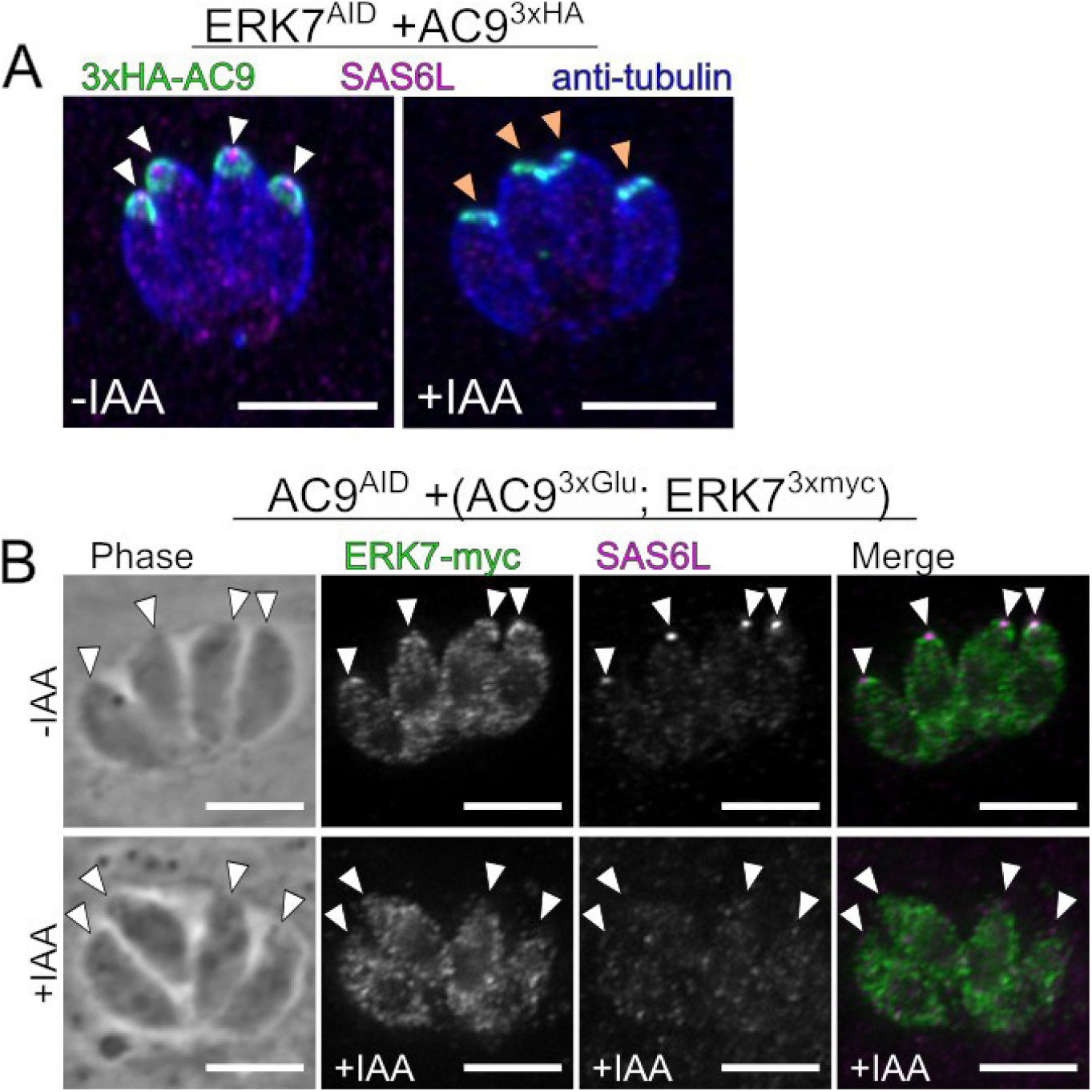
(A) AC9 localization is unaffected by ERK7 degradation. ERK7^AID^(3xHA-AC9) parasites were grown in the presence or absence of IAA and stained with anti-HA (AC9; green), anti-SAS6L (magenta), and anti-β-tubulin (blue). Note anti-β-tubulin does not stain conoid, likely due to antigen accessibility. Images are maximum intensity projects of confocal stacks. White arrowheads indicate SAS6L-positive conoid puncta. Orange arrowheads indicate expected localization of missing SAS6L puncta in ERK7^AID/IAA^ parasites. (B) AC9^AID^(+AC9^3xGlu^; ERK7-3xmyc) parasites were grown in the presence or absence of IAA and stained with anti-myc (ERK7; green) and anti-SAS6L (magenta), a marker for the parasite conoid. Arrows indicate the parasite apical end. Note the bright SAS6L foci in the -IAA parasites that are lost in +IAA. All scale bars: 5 μm.

**Table S1.** Data collection and refinement statistics. Statistics for the highest-resolution shell are shown in parentheses.

| Data collection |  | Refinement |  |
| --- | --- | --- | --- |
| Wavelength | 1.541 | Reflections used in refinement | 39043 (3083) |
| Resolution range | 29.89 - 2.1 (2.175 - 2.1) | Reflections used for R-free | 1718 (141) |
| Space group | P 1 | R-work | 0.1583 (0.1787) |
| Unit cell | 50.8897 52.9379 67.8003<br>82.55 84.5785 81.35 | R-free | 0.2119 (0.2329) |
| Total reflections | 93549 (4051) | CC(work) | 0.961 (0.919) |
| Unique reflections | 39049 (3083) | CC(free) | 0.924 (0.809) |
| Multiplicity | 2.4 (1.3) | Number of non-hydrogen atoms | 5986 |
| Completeness (%) | 96.68 (76.39) | macromolecules | 5468 |
| Mean I/sigma(I) | 15.17 (3.48) | ligands | 20 |
| Wilson B-factor | 17.81 | solvent | 498 |
| R-merge | 0.05342 (0.1976) | Protein residues | 682 |
| R-meas | 0.0665 (0.2744) | RMS(bonds) | 0.004 |
| R-pim | 0.03903 (0.1898) | RMS(angles) | 0.61 |
| CC1/2 | 0.997 (0.894) | Ramachandran favored (%) | 98.33 |
| CC* | 0.999 (0.972) | Ramachandran allowed (%) | 1.67 |
|  |  | Ramachandran outliers (%) | 0.00 |
|  |  | Rotamer outliers (%) | 0.17 |
|  |  | Clashscore | 1.45 |
|  |  | Average B-factor | 29.43 |
|  |  | macromolecules | 29.18 |
|  |  | ligands | 44.08 |
|  |  | solvent | 31.59 |
|  |  | Number of TLS groups | 16 |

**Figure S6.**
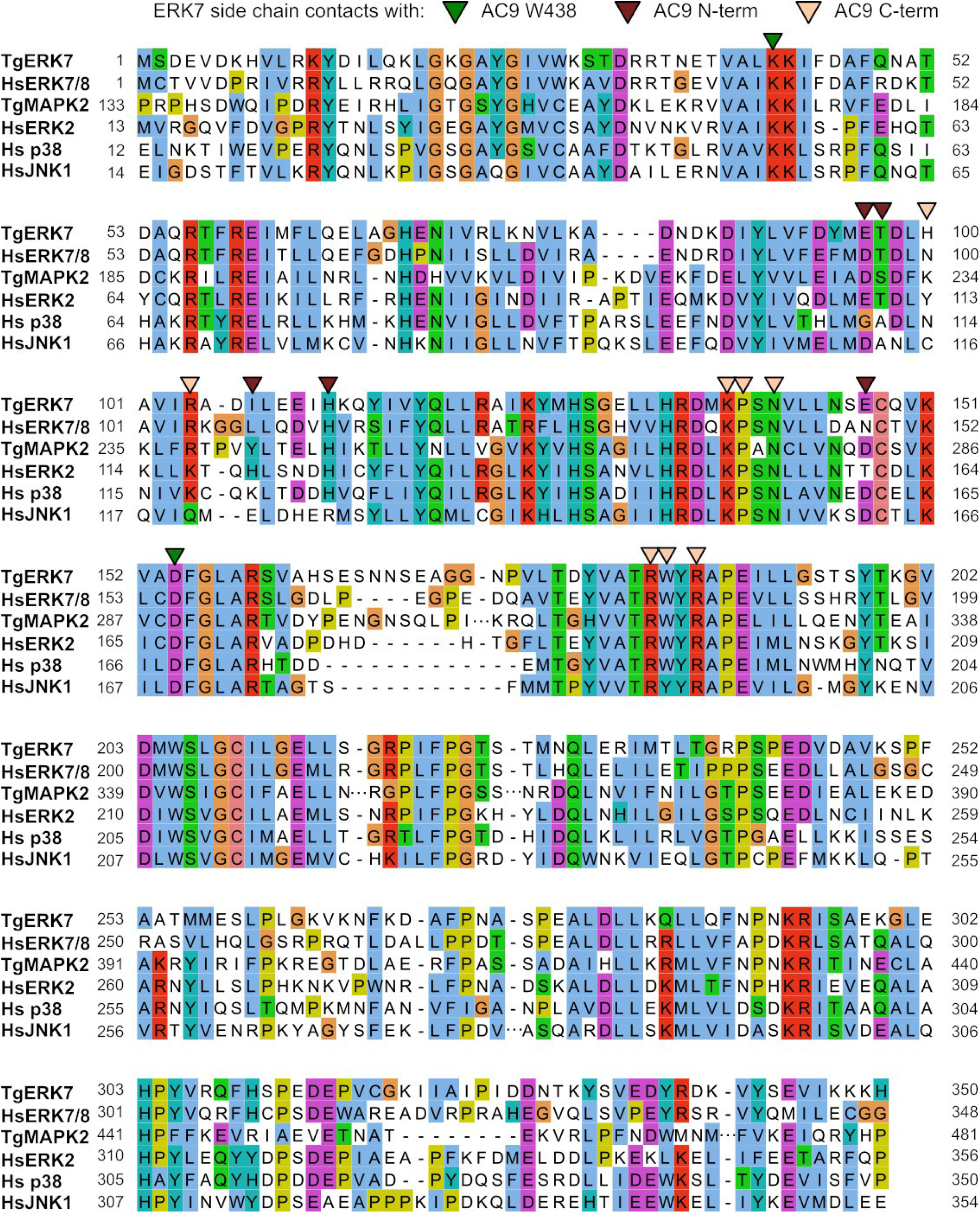
Alignment of TgERK7 and TgMAPK2 kinase domains with human MAPKs. TgERK7 side chains that make contact with AC9_419-452_ in our crystal structure are marked as indicated. Large gaps were removed from alignment and are indicated with ellipses.

**Figure S7.**
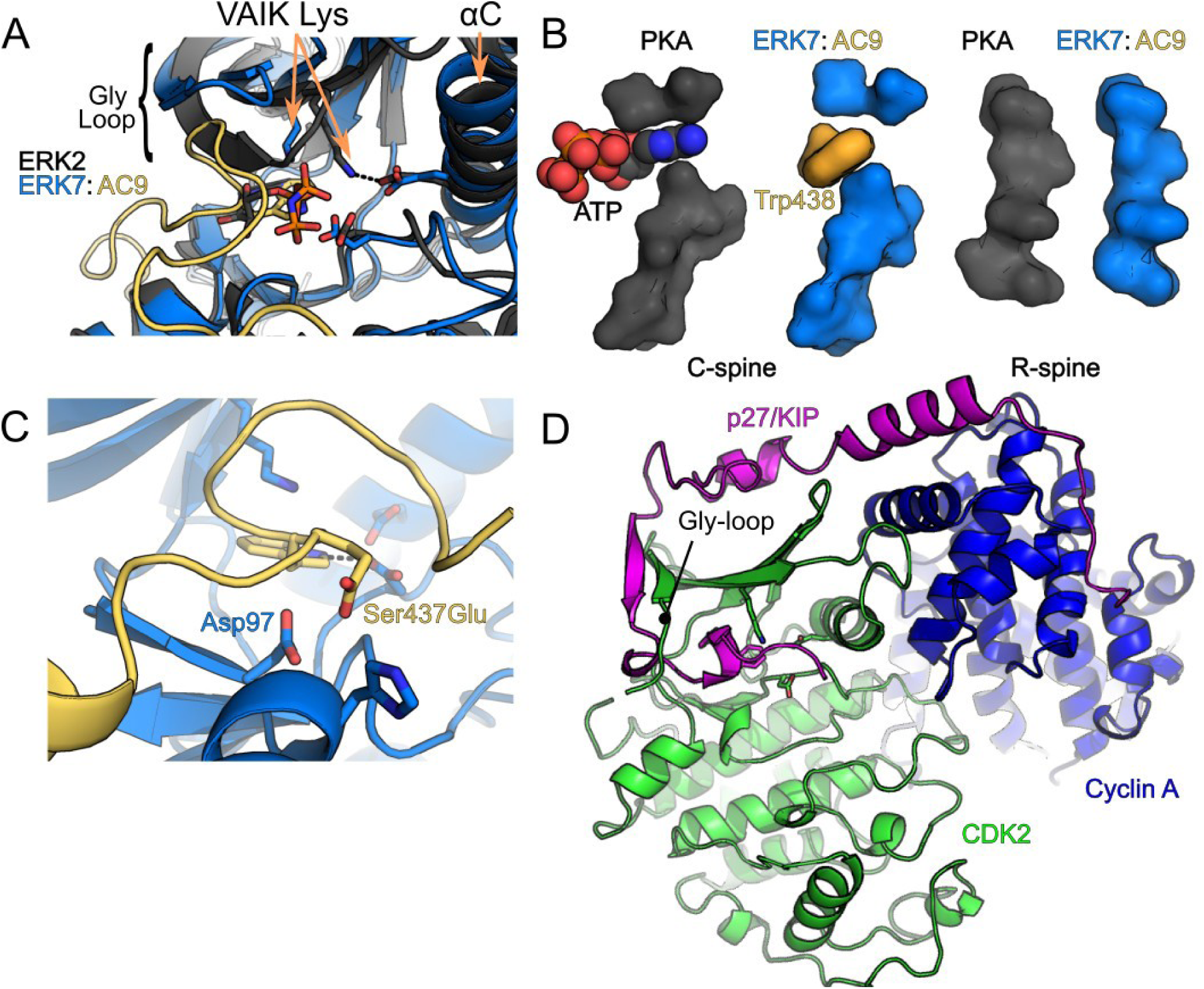
Comparison of ERK7:AC9 with other kinase structures. (A) The structure of activated, AMP-PNP-bound ERK2 (black; 6OPG) is overlaid with ERK7:AC9 (blue and yellow) and the Gly-loop, catalytic lysine, and αC helices are indicated. Note that the ERK7:AC9 Gly-loop is held in an inactive conformation by AC9, which keeps the αC-Glu from salt bridging with the VAIK Lys.(B) The “C-spines” (PKA: V57, A70, M128, L172, L173, I174, L227, M231; TgERK7: V27, A40, L99, V142, L143, L144, I210, L214) and “R-spines” (PKA: L95, L106, Y164, F185; TgERK7: L64, L76, H134, F155) of activated, ATP-bound PKA (black; 1ATP) is shown in comparison to ERK7:AC9 (blue and yellow). While the R-spine is intact, the C-spine is not fully complete by Trp438; ERK7 is not in an active conformation when bound to AC9, as the Gly-loop is in an open conformation, and the VAIK:α-C Glu salt bridge are not formed (see (A) and Figure 5). In addition, AC9 displaces the ERK7 activation loop (see Figure 5D,E). (C) Mutation of AC9 Ser437 to Glu would clash with the side chain of Asp98 (<3.5 Å between the carboxylate side chains) when AC9 is in the optimal conformation for binding ERK7. (D) Overview of the CDK2:p27/KIP:Cyclin inhibitory complex (1JSU). p27 (magenta) wraps around the cyclin A (blue) and CDK2 (green), unfolding the CDK2 Gly-loop and inserting it’s C-terminal residues into the active site, replacing nucleotide.

## Notes

#### Summary of Updates

Additional biochemical data added (Figures 5, 6, S3)

